# Ubiquity of inverted ’gelatinous’ ecosystem pyramids in the global ocean

**DOI:** 10.1101/2024.02.09.579612

**Authors:** Lombard Fabien, Guidi Lionel, Manoela C. Brandão, Coelho Luis Pedro, Colin Sébastien, Dolan John Richard, Elineau Amanda, Josep M Gasol, Grondin Pierre Luc, Henry Nicolas, Federico M Ibarbalz, Jalabert Laëtitia, Loreau Michel, Martini Séverinne, Mériguet Zoé, Picheral Marc, Juan José Pierella Karlusich, Rainer Pepperkok, Romagnan Jean-Baptiste, Zinger Lucie, Tara Oceans Coordinators, Stemmann Lars, Silvia G Acinas, Karp-Boss Lee, Boss Emmanuel, Matthew B. Sullivan, Colomban de Vargas, Bowler Chris, Karsenti Eric, Gorsky Gabriel

**Author notes:** A list of authors and their affiliations appears at the end of the paper.

## Abstract

Plankton are essential in marine ecosystems. However, our knowledge of overall community structure is sparse due to inconsistent sampling across their very large organismal size range. Here we use diverse imaging methods to establish complete plankton inventories of organisms spanning five orders of magnitude in size. Plankton community size and trophic structure variation validate a long-held theoretical link between organism size-spectra and ecosystem trophic structures. We found that predator/grazer biomass and biovolume unexpectedly exceed that of primary producers at most (55%) locations, likely due to our better quantification of gelatinous organisms. Bottom-heavy ecosystems (the norm on land) appear to be rare in the ocean. Collectively, gelatinous organisms represent 30% of the total biovolume (8-9% of carbon) of marine plankton communities from tropical to polar ecosystems. Communities can be split into three extreme typologies: diatom/copepod-dominated in eutrophic blooms, rhizarian/chaetognath-dominated in oligotrophic tropical oceans, and gelatinous-dominated elsewhere. While plankton taxonomic composition changes with latitude, functional and trophic structures mostly depend on the amount of prey available for each trophic level. Given future projections of oligotrophication of marine ecosystems, our findings suggest that rhizarian and gelatinous organisms will increasingly dominate the apex position of planktonic ecosystems, leading to significant changes in the ocean’s carbon cycle.

---

Marine plankton drift with ocean currents, with hundreds of thousands of species from metazoans to prokaryotes, as well as viruses^1–3^. Together, plankton constitute the base of pelagic food webs and modulate global biogeochemistry^4^. Understanding the mechanisms underpinning plankton ecosystem structure is a major focus in planetary ecology^5,6^, however most studies have reported fragmented views of plankton communities partitioned by size ^2,7,8^ or taxonomic group^9^ mostly because of sampling and analysis limitations. A global and inclusive view of the full trophic organization of whole plankton communities is lacking, hampering our understanding of trophic equilibria and dynamics in marine ecosystems.

A conventional technique for holistically assessing ecosystem structure and function is the ‘size spectra approach’, generalising Elton’s pyramid^10^ of numbers that describes the inverse relationship between the size of organisms and their abundance. The Elton pyramid has been reformulated into biomass^11^ and trophic pyramids^12^ as well as biomass or biovolume size spectra (BSS^13,14^) and normalised biomass or biovolume size spectra (NBSS^15^). In the plankton, primary producers are small in size and consumed by grazers and predators of increasing size with trophic level^16^. Because of this principle, the slope of the continuously decreasing NBSS (S_NBSS_) is linked to the balance between consumers and prey^17^ and represents a proxy for the slope of the trophic pyramid (S_Trophic_). Thus, S_NBSS_ < -1 is assumed to represent conventional ‘bottom-heavy’ trophic pyramids (*i.e.*, S_Trophic_ < 0), while flatter slopes are associated with ‘top-heavy’’ inverted pyramids (S_Trophic_ > 0; Fig. 1b). S_NBSS_ is an essential input to numerous theories and models of community metabolism, energy use and transfer efficiency, which attempt to uncover the fundamental mechanisms underlying trophic relationships^18^. The size-spectra approach integrates the full size range of organisms as a single ecological object, yet only few studies have used it over the wide range of plankton^19–25^ and none have done so at global spatial scale crosslinked with taxonomic or functional properties (aside from modelling exercises^26^).

**Figure 1.**
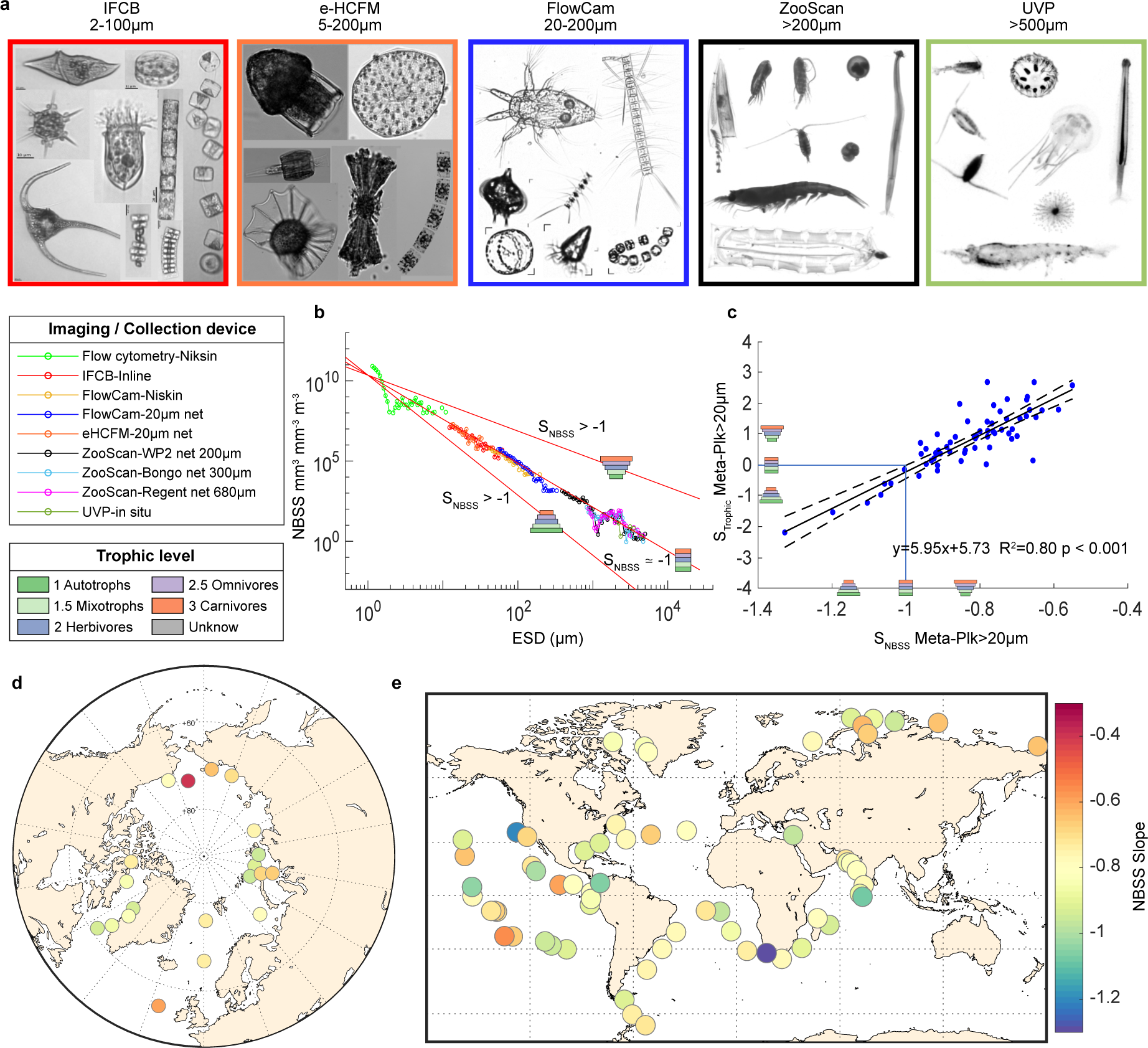
: *Tara* Oceans multi-imaging framework to assess the trophic structures of open ocean plankton ecosystems at global scale. a) Examples of images obtained with the different quantitative imaging devices (e- HCFM - environmental High Content Fluorescence Microscopy, IFCB - Imaging FlowCytoBot, FlowCam, Zooscan and Underwater Vision Profiler (UVP) ; for complete image collection including scale bars see Table S1. b) Full normalized biovolume size spectra (NBSS) from station 173, reconstructed by combining the different size classes, and plotted as a function of organism size (ESD: equivalent spherical diameter). Theoretical links between NBSS slopes (S_NBSS_) and trophic pyramid structure (S_Trophic_) are also indicated. c) Observed relationship between S_NBSS_ and S_Trophic_ for the entire meta-plankton community >20µm (Meta- Plk >20µm) across all *Tara* Oceans samples . d) S_NBSS_ for Arctic (Meta-Plk >0.8 µm) and e) world ocean (Meta- Plk >20µm) plankton ecosystems

According to theory^10^, energy loss between trophic levels should lead to ‘bottom-heavy’ pyramids with higher biomass of primary producers than consumers. While this pattern is consistently observed in terrestrial food webs, some rare observations or models^27^ ^and^ ^references^ ^therein^ have suggested that marine food webs may instead be structured as inverted pyramids^9,28^. However, because of energetic and predator-prey size constraints these inverted pyramid structures have been interpreted as being a result of sampling artefacts^16^. Even if inverted pyramids may result from high turnover rates of producers compared to consumers^27^, the mechanisms generating bottom-heavy versus inverted top-heavy pyramids remain unclear, together with their consequences on energy transfer in ecosystems.

Here we integrated six optical and imaging technologies deployed during the *Tara* Oceans (TO; 2009- 2012) and *Tara* Oceans Polar Circle (TOPC; 2013) expeditions to examine variation in plankton community structure across global taxonomic and spatial scales (Fig. 1a,b). We used multiple complementary sampling strategies and devices (inline pumping systems, Niskin bottles, peristaltic pumps, plankton nets) with diverse quantitative optical/imaging instruments including flow cytometry, Imaging Flow Cytobot (IFCB), environmental High-Content Fluorescent Microscopy (eHCFM), Flowcam, Zooscan, and Underwater Vision Profiler (UVP) (Fig. 1a; see Methods) to estimate the concentration of plankton across 5 and 15 orders of magnitude in size and biovolume, respectively. The imaged organisms were sized, sorted taxonomically using semi-automated image classification^29^, and aggregated into community-relevant ecological groups related to function, abundance and/or trophic level (*i.e*., primary producer, mixotroph, herbivore, omnivore, carnivore; see Methods). Per-organism size measurements were used to compute NBSS for each instrument at each sampling site, and data were then combined to obtain a global scale, homogeneous quantification of plankton, hereafter called ‘metaplankton’ (Fig. 1b). The reconstructed metaplankton communities are composed of organisms ranging in size from 0.8 µm to several cm for the Arctic Ocean (Meta-Plk >0.8 µm, Fig 1d), and from 20 µm to several cm (Meta-Plk >20 µm, Fig 1e) in the rest of the global ocean depending on the variety of measurements done (Fig. S1). For each metaplankton assemblage, we calculated the S_NBSS_ and S_Trophic_ (see Methods). Sampling occurred mostly during day time but night observations are available for cross comparison. The results were further compared to 18S rDNA metabarcoding data from the same sites, and metaplankton products were finally converted to carbon units to assess their ecological and biogeochemical relevance.

We found strong correlations between the slopes extracted from the two indicators (S_NBSS_ and S_Trophic_) of ecosystem size-spectra and trophic structures (Fig. 1c, Fig. S2), confirming for the first time the theoretical link between them. Since S_NBSS_ is independent of taxonomy and trophic level, this result provides strong support for our taxonomic and trophic assignment of organisms (Table S4), and indicates that both proxies of community trophic structure are consistent and interchangeable. Overall, a predominance of top-heavy, inverted trophic community structures was found at the global scale (68% and 74% based on biovolume S_Trophic_ and S_NBSS_, respectively; Fig. 2a). Focusing on the 20 Arctic stations, 45% and 75% of analysed communities were top-heavy based on S_Trophic_ or S_NBSS_, respectively (Fig. 2a). This top-heavy trophic structure of marine plankton was robust and consistently found regardless of the particular dataset, including when using carbon biomass conversions, adding bacteria and pico-nano plankton counts from FACScalibur flow-cytometry measurements, or comparing to trophic assessments based on taxonomic annotation from DNA metabarcoding data^2,7^ (Fig.2, Fig. S3-5), with correlation between S_NBSS_ and S_Trophic_ remaining valid (Fig. S2). This top-heavy structure is even reinforced when based only on night observations, when migrant zooplanktonic grazers and predator migrate to the ocean surface (Fig. S5). Bottom-heavy ecosystems (the norm on land) were relatively rare (4% and 11% at global scale when assessed with S_NBSS_ and S_Trophic_, respectively, and only 0 and 30% in the Arctic Ocean). They appear limited to relatively productive conditions (Fig. 1d,e, S4) from coastal upwelling (e.g., Benguela, Panama, and California upwelling systems at, respectively, TO-Stations 67, 140, and 133), or phytoplankton blooms such as occurring at the sea ice margin (*e.g*., TOPC-Stations 173, 175 and 188).

**Figure 2:**
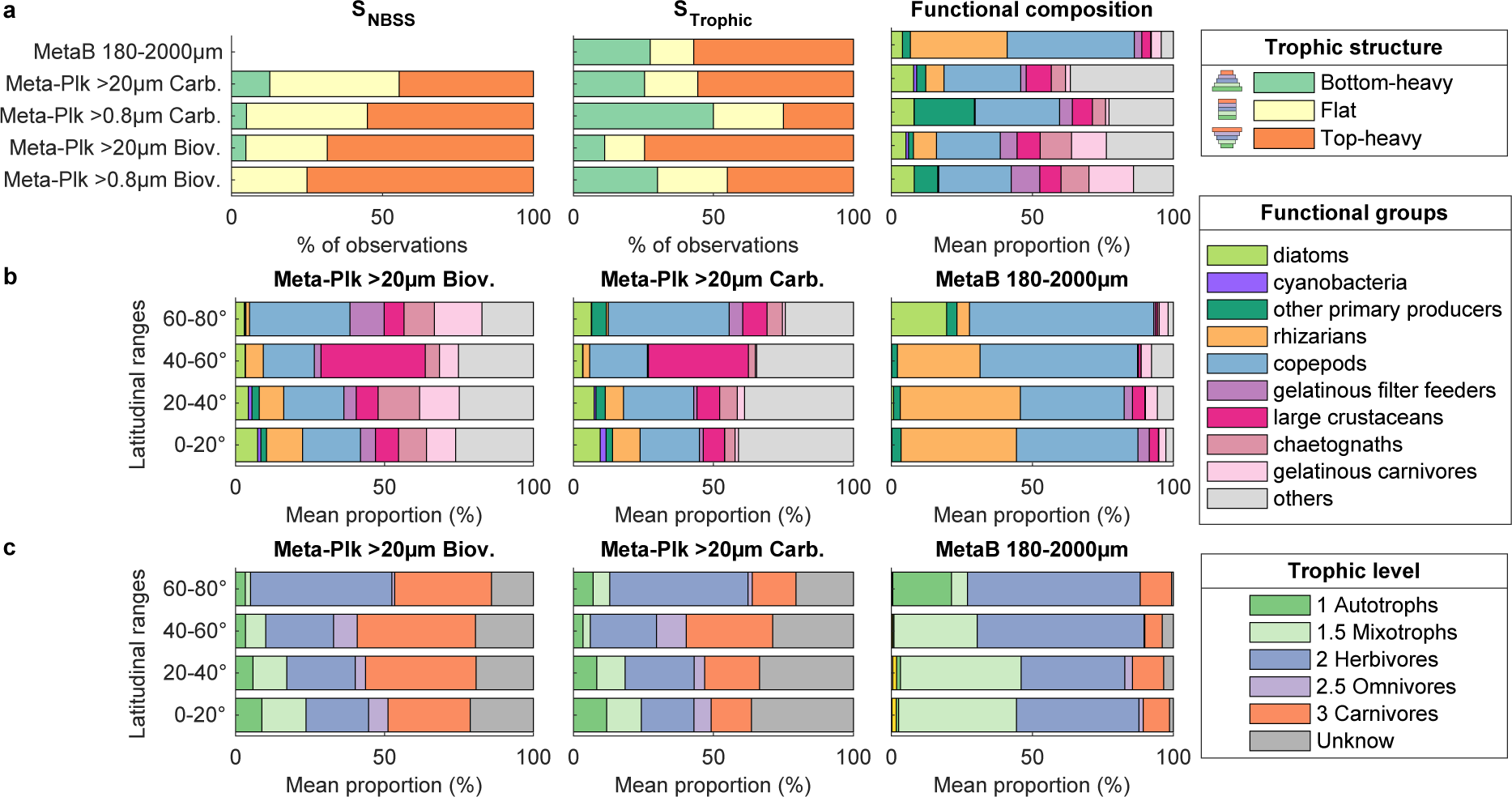
Global predominance of top-heavy trophic pyramids in the world marine plankton. a) The proportion of bottom-heavy, flat and top-heavy trophic community structures was established on the basis of NBSS (S_NBSS_ <-1,1 ; -1.1-0.9 ; >-0.9) and Trophic (S_Trophic_ < -0.25, -0.25-0.25 and >0.25) slopes. These were calculated for metaplankton datasets from the Arctic (Meta-Plk >0.8µm) and global (Meta-Plk >20µm) Oceans, either using biovolume (Biov.) or carbon biomass (Carb.) for computation. The mean metaplankton functional composition of each dataset was also extracted. **b)** Latitudinal variations of the different functional and plankton groups (c) trophic levels calculated for the global ocean metaplanktonic datasets (Meta-Plk >20µm;) using either biovolume (Biov.) or carbon biomass (Carb.). Independent calculations from DNA metabarcoding datasets from the meso-planktonic size fraction were also included.

This high proportion of top-heavy trophic structures originates from the relative proportions of certain planktonic functional groups. The Arctic metaplankton community (Meta-Plk >0.8µm, Fig. 2a, Table S1) is composed of a high proportion of gelatinous organisms (including carnivorous chaetognaths, gelatinous predators, and gelatinous herbivorous filter feeders, 35% of the total biovolume), copepods (25%), large crustaceans (7%), diatoms (8%), and other phytoplankton (8%). This result contrasts with the classical paradigm of the Arctic plankton food web as being strongly dominated by diatoms, copepods and krill^49–51^, and could result from our holistic approach which associates classical data from nets with non-destructive *in-situ* image acquisition. Furthermore, the poor preservation of gelatinous zooplankton in formaldehyde^30^, such as ctenophores known to be important in Arctic ecosystems, could have led to the underestimation of the predominance of gelatinous organisms in previous studies.

A similar functional compositional pattern in plankton community structure was observed throughout the global ocean (Meta-Plk >20µm, Fig. 2a), with gelatinous organisms (filter-feeding tunicates and carnivores including cnidarians, ctenophores and chaetognaths), copepods, large crustaceans, diatoms, other phytoplankton, and rhizarians representing approximately 29%, 22%, 8%, 5%, 2% and 8% of the total biovolume, respectively. Global predominance of gelatinous plankton is unexpected since they typically represent a small fraction in previous global plankton estimates^31–33^. Conversion from biovolume to carbon biomass decreases this contribution to 10% (Fig. 2b; Fig. S3, S4c,d), which is still an order of magnitude greater than previously reported values (<1%^34^), suggesting that former studies largely underestimated the content of fragile gelatinous organisms^35,36^. Though an imperfect quantitative metric^37^, metabarcoding data are in agreement with these image-based organismal abundances, albeit with noticeable deviations for copepods and rhizarians.

NBSS and trophic slopes show no latitudinal trends (Fig. 1d,e, S3, S4), in agreement with results of a theoretical modelling framework^38^. Small differences between polar and tropical environments only appear when looking at plankton functional groups or trophic levels (Fig. 2b, c). Surprisingly, the Arctic food web does not strongly differ from the global ocean in terms of functional and trophic structures, other than an increase in the abundance of rhizarian or mixotrophs in tropical zones, and of copepods in the Aarctic ecosystems (Fig. 2b, c). While Arctic ecosystems are less diverse^7^ and structurally simpler^39^, our results suggest that energy transfer through Arctic plankton food webs follow the same principles as elsewhere in the global ocean at the trophic and functional levels. We further investigated this unexpected similarity by examining the relationship between the biovolumes of consumers (herbivorous and carnivorous) and their prey (Fig. 3). Our findings indicate that the biovolume of consumers depends on that of their prey, yet with no direct proportionality. Rather the biovolume of predators often exceeds that of prey in low food conditions, and prey biovolume exceeds that of predators only in exceptionally prey-rich environments, thereby generating classical terrestrial-like pyramids, in line with previous observations in both terrestrial^40^ and oceanic ecosystems^9,41^, and even between viruses and their bacterial prey^42^. This emergent property of plankton trophic structure holds true from Arctic to global ocean ecosystems, and across trophic levels, from primary producers to carnivores (Fig. 3). This indicates that the trophic structure of ecosystems is primarily driven by prey stocks, and not necessarily by their productivity, as also confirmed by the lack of correlation between trophic structure indices and satellite-derived proxies of ecosystem productivity or chlorophyll (Fig. 4a, b).

**Figure 3:**
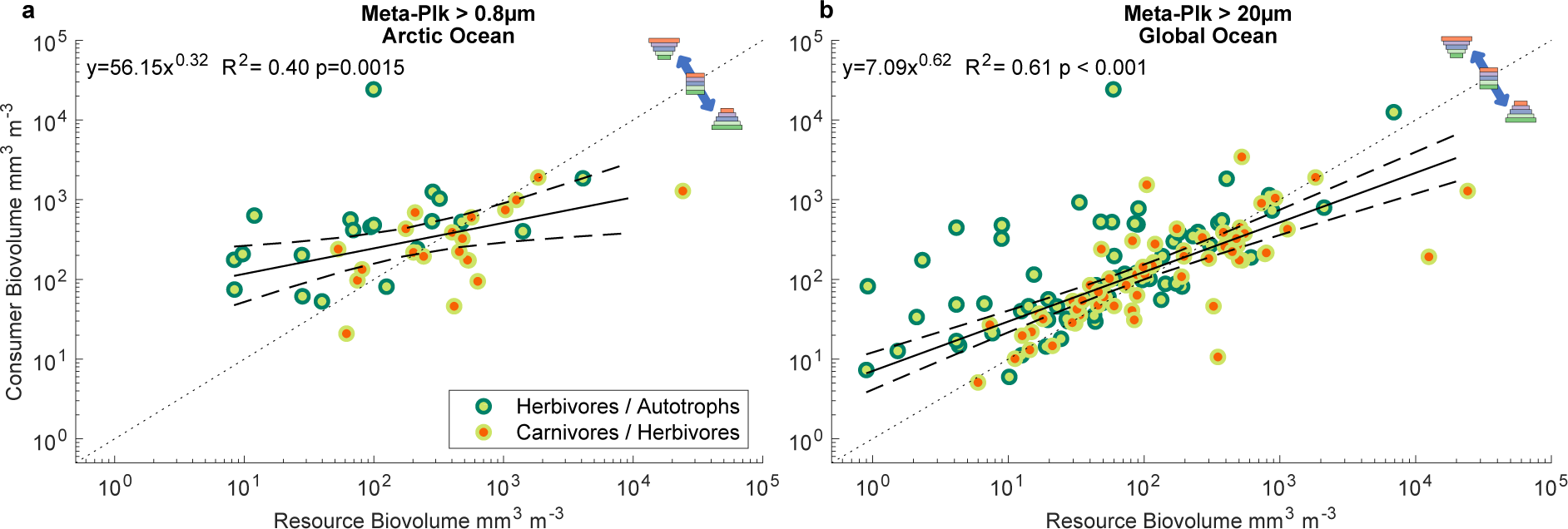
Biovolume relationships between prey organisms and their predator for the Arctic and global Oceans. Prey versus predator biovolume relationships are shown separately for the autotrophs/herbivores (deep/light green dots) and herbivores/carnivorous (light green/red dots) couples. Deviation from the 1:1 relationships are indicators of trophic structure.

**Figure 4:**
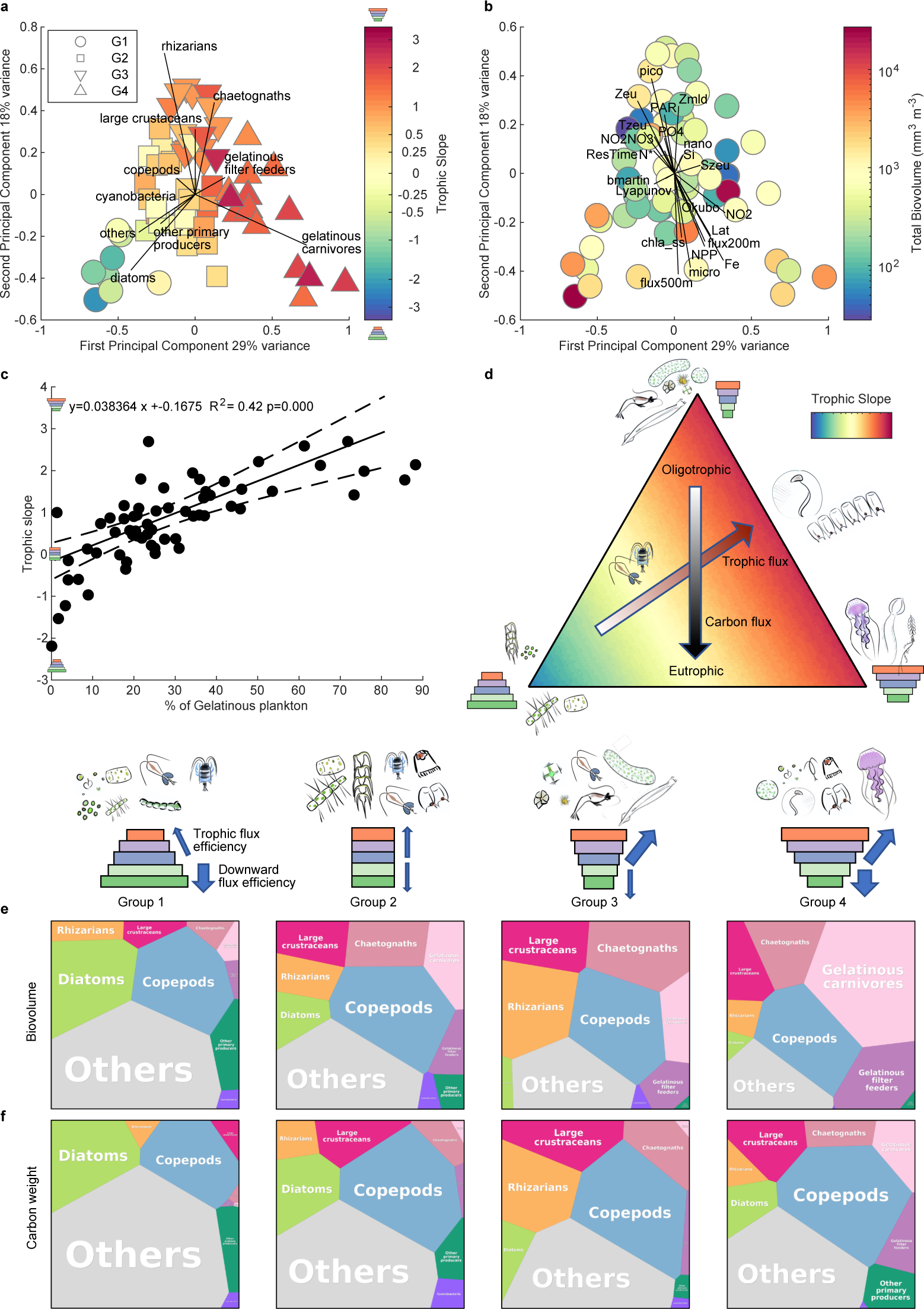
**Link between ecosystem functional composition, trophic structure, and environmental properties**. a) Principal Component Analysis (PCA) performed on the main taxo-functional groups from *Meta- Plk >20µm*, and with correlation of the functional groups with PCA components (arrows) and their associated trophic slopes (color scale). Four groups of stations displaying different trophic and functional signatures could be detected using Euclidean distances and Ward linkage clustering on PCA coordinates: Group 1 (circles), 2 (squares), 3 (inverted triangles), 4 (triangles), respectively dominated by (1) diatom and phytoplankton; (2) copepod; (3) rhizarian, chaetognath and large crustacean and (4) gelatinous plankton. b) The same PCA analysis with correlation of environmental properties (see methods for details on the contextual parameters integrated in this analysis) with PCA axes while the colour scale represents total biovolume. c) Relationships between the trophic slopes and the percentage of gelatinous plankton (gelatinous carnivores + filter feeders + chaetognaths). d) Conceptual scheme of the different ecosystem states observed in terms of trophic and functional structure, together with their potential links with the carbon flux in the water column or in the trophic chain. e) Observed taxo-functional groups’ average biovolume (e) and carbon weight (f) proportions for each group of stations defined in (a) and corresponding to four ecosystem states. See S6 for the effect of adding microbes to these structures. Visualisation obtained from http://bionic-vis.biologie.uni-greifswald.de/.

To investigate whether the functional composition of metaplankton communities is associated with specific environmental conditions, we conducted a Principal Component Analysis (PCA; Fig. 4a) on which correlations with environmental features were projected (Fig. 4b). Three extreme types of functional composition emerged, with a direct link to trophic slopes (Fig. 4a and d). The bottom-heavy communities were associated with low principal component values on axis 1 and 2, while the two other extremes were strongly top-heavy and characterised by high proportion of gelatinous organisms (Fig. 4d). Most environmental features are associated with axis 2 characterising the eutrophic (negative values, high biovolumes correlated with NPP, chlorophyll *a,* carbon flux and iron concentration) to oligotrophic gradient (positive values, correlated with greater depth of mixed layer Z_mld_ euphotic zone Z_eu_ and PAR among others). While the trophic slope is strongly related to axis 1, a few environmental parameters are weakly associated with it (Martin’s *b*, and S_Zeu_), further establishing the trophic structure as an ecosystem emergent property that is largely independent of environmental forcing (Fig. 4b) but significantly associated with the percentage of gelatinous organisms (Fig. 4c).

The three extremes in plankton taxo-functional composition could be separated into 4 groups of sampling sites (Fig. 4a, Table S2) for which the main biotic composition was assessed (Fig. 4e). Group 1 has a high representation of diatoms (18% biovolume as a mean) and low biovolume of gelatinous organisms (4.7%), and is observed in coastal and equatorial (Pacific) upwelling zones (Fig. S6a). Group 2 is characterised by flat trophic structures and comprises a relatively equilibrated taxo-functional composition (still including 21% of gelatinous organisms). Group 3 is characterised by a higher rhizarian (16%) and chaetognath (17%, total gelatinous ornanisms at 29%) biovolume composition, and is observed mostly in tropical oligotrophic regions. This is consistent with previous findings of high biomass of rhizarians, often having adaptations convergent with gelatinous organisms^43^ in oligotrophic gyres^44^. Finally, Group 4 has a large proportion of gelatinous organisms (55% biovolume), both in the form of filter feeders (12%, e.g., salps appendicularians) and carnivores (32%, e.g., jellyfishes, ctenophores, chaetognathes), and is observed in coastal areas.

Using carbon biomass (Fig. S6b) or adding flow cytometry data to include bacteria and pico- nano- plankton in the analysis (Fig. S6c) does not alter our results but decreases the proportion of gelatinous organisms (e.g., to 5% in Group 2; 16% in Group 4) and increases by a constant proportion the heterotrophic bacteria (2.5-8.2% biovolume) and cyanobacteria (3.8-7.1%) pools. Likewise, copepods represent a constant 19-25% biovolume in every ecosystem state (Fig. 4e, S6b). Our findings suggest that, although copepods and large crustaceans both have carnivorous representatives, the predominance of top-heavy trophic structures observed at global scale is directly connected with gelatinous organisms (Fig. 4c,d). More importantly, top-heavy trophic structures are both observed in oligotrophic (Group 3) and eutrophic conditions (Group 4), suggesting that other intrinsic ecosystem properties are responsible for such observations.

When previously observed, top-heavy ecosystem structures were believed to result from specific biological and ecosystem properties^27,45,46^. These properties are however commonly met for planktonic ecosystems especially when considering gelatinous plankton. Plankton turnover rates are high for autotrophs with time scale of growth in the range of hours to days^47^ while their grazers and predators have life cycles ranging from a few days to months^48^. Gelatinous plankton are known to have relatively low metabolic expenses compared to their feeding capacities^49^, therefore increasing efficiency of energy transfer to higher trophic levels. They are also able to forage on prey that are several orders of magnitude smaller than those of similar sized predators^50^, therefore short-circuiting food web structures but probably providing lower food quality to higher trophic levels^51^. Finally, gelatinous plankton also have the capacity to consume their own biomass and shrink to survive over long starvation periods^52^, further increasing the life span difference between predator and prey. It should also be noted that the overall variability in the biovolume of autotrophs is larger than that of consumers or predators (Fig 3), implying that predators (more stable) have a larger resilience and buffering capacity against seasonal variations than their prey (more variable with intense bloom and bust cycles). All of the above favours the emergence of top-heavy ecosystem structures in the ocean.

In conclusion, our results show that top-heavy planktonic ecosystems are observed worldwide in the upper ocean, except in specific conditions (early blooms) when and where the decoupling between predator and prey is largest. They are associated with a dominance of gelatinous organisms (>55% of observations) in both polar and non-polar regions. Inverted-gelatinous dominated ecosystems are associated with oligotrophy and late blooms, but not with latitudinal gradients, while classical, terrestrial-like pyramids are associated with early bloom eutrophic conditions. Most open ocean plankton ecosystems appear to be organised along three extreme communities: diatom/copepod dominated (early bloom eutrophy), rhizarian/chaetognath dominated (warm water oligotrophy), and gelatinous dominated (late bloom eutrophy). The observed plankton ecosystem structures have consequences for biogeochemical fluxes (Fig 4b, d). Eutrophic systems dominated by diatoms and copepods transport a higher proportion of new production to depth, but with lower trophic transfer efficiency. Oligotrophic rhizarian systems, on the other hand, exhibit higher transfer efficiency but lower vertical export. Systems dominated by gelatinous organisms are associated with both high vertical flux^53^ and high trophic transfer. Current climate change projections highlight the possible ‘tropicalisation’ of the marine environment^54^, *i.e.*, an increase of stratification and oligotrophy^55^. Our results suggest that this will lead to increased rhizarian and gelatinous plankton-based ecosystems in the ocean.

## Methods

### Sampling

The complete sampling protocols used in *Tara* Oceans and *Tara* Oceans Polar Circle expeditions are detailed in^56^. In order to compare as many measurements as possible, we focused on samples collected from Niskin bottles in the surface layer (0-3m), the ship’s inline water intake located 2 m below sea surface, and plankton nets deployed at various depths with mesh size of 5µm (0-5m), 20µm (0-5m), 200µm (“WP2 net”, 0-100m), 300µm (“bongo net”; 0-500m), and 680µm with silk mesh (“Regent net”; 0-500m). We also used the Underwater Vision Profiler^57^ mounted on the Rosette which recorded *in-situ* images of > 600 µm plankton. For nets and UVP, we only used day-time samples (defined as when the sun azimuth was above the horizon with a 2° margin to incorporate dusk conditions) and samples collected in the upper 200m of the water column. We choose to not include night samples to keep a conservative bias (potential underestimation of grazers and predators). Similar results were obtained when substituting night samples to day samples when available (Fig S3, S5), and top-heaviness is even more present, a pattern coherent with vertical migrations of herbivorous and carnivorous organisms.

A full set of optical or imaging devices were used to count, qualify and measure plankton. For the full *Tara* Oceans cruise (stations up to 154), the different set used includes 1) cells counts using a FACSalibur flow cytometer 2) environmental High Content Fluorescence Microscopy^68^ (e-HCFM) using sample originating from a 5 and 20µm mesh size nets, 3) samples collected with different nets and imaged with the Zooscan^58^ and 4) in situ observations done with the Underwater Vision Profiler^57^ (UVP-5). For the *Tara* Oceans Polar Circle, this sampling scheme was complemented by on-board instruments including 5) Accuri flow cytometer, 6) Imaging FlowCytoBot (IFCB^59,60^) and 7) FlowCam analyzer^61^ (See Supplementary Information for further details on sampling).

### Ecotaxa processing and post-processing

Images from different sources described above were identified by taxonomic experts using the online software Ecotaxa^62^. The remaining images were predicted in Evotaxa by machine learning methods. The different Ecotaxa projects with their total number, percentage of validated objects and the link to them are given in Table S3.

Depending on the data source inspected, the completion of validation varied (Table S3, Fig. S1), but was complete for organisms of larger fractions (WP2, Bongo, Regent, UVP) and within the Arctic, when numerous instruments were deployed simultaneously. Finally, 22,309 and 25,095 images were identified on the eHCFM 5µm and eHFCM-20µm datasets, with a reasonable prediction of the rest of the dataset^63^. In eHFCM-20µm, due to sample preparation, a large number of images corresponds to multiple organisms overlapping each other which explains the large biovolume of “other unidentified’’ organisms in this dataset. An extensive quality check of metadata (volumes of water, volume of sample inspected) was conducted.

All results from Ecotaxa were extracted as individual text files. Taxonomic annotation, morphometric measurements and essential metadata (volume of water collected, volume of sample inspected) were used to calculate the biovolume (in mm^3^) of each particle collected (plain area biovolume, extruded area biovolume and ellipsoidal equivalent biovolume assuming prolate ellipsoids). While none of these biovolumes gives perfect results, we choose to use the ellipsoid biovolume for every instrument.

Organisms abundances (ind. m^-3^) and biovolumes (mm^3^ m^-3^) were calculated for each taxonomic annotation but also with several levels of regrouping: 1) total, 2) living or non-living 3) a functional annotation and 4) a trophic annotation. We chose to define 23 planktonic functional groups corresponding to broad taxonomic groups with important ecological functions (e.g. ^64^). After a preliminary analysis, low abundance groups with similar functions were grouped mostly under the label “other primary producers” (for autotrophic/mixotrophic groups) and “others”. Trophic annotations corresponding to each taxon were used to regroup autotroph taxa as trophic level 1, mixotrophs (1.5), grazers (2), omnivorous (2.5) and carnivorous (3) based on bibliographic research (e.g.^65–68)^ as well as consultations with taxonomic experts. Non-living or non-feeding were attributed to the trophic level -1. For uncertain cases, we followed a conservative approach and allocate the status of grazers (2) for any heterotroph having a non-strict omnivorous or carnivorous behaviour, notably concerning copepod species in which prey switching may occur or following the recommendation of Flynn et al^69^ for microplankton organisms. Any organisms for which the trophic mode could not be attributed were kept as undetermined (noted 3.5). The full list of functional and trophic annotations linked with their Ecotaxa taxonomic label can be found in Table S4.

### Normalized biovolume Size Spectra (NBSS) calculation

Following^15^, biovolume size spectra (BSS) and normalized biovolume size spectra (NBSS) were obtained under a harmonic scale of biovolume starting from 10^-12^ to 10^4^ mm^3^ with biovolume size-class increasing exponentially with minimal and maximal biovolumes of each size-class (Bv_min_ and Bv_max_) defined such as Bv_max_= 2^0.25^ Bv_min_. BSS was obtained by summing the biovolume of each object belonging to each size-class while NBSS was obtained by dividing BSS by the biovolume width of each size-class (i.e. Bv_range_=Bv_max_- Bv_min_). BSS and NBSS spectra were calculated for initial taxonomical identity and for each level of regrouping (i.e. functional type and trophic level).

The BSS is roughly comparable to a pyramid of biomass^11^ while the NBSS is representative of pyramids of numbers^70^ with a scaling factor of Bvmean/Bvrange to recover counts within a size range^71^.

### Metaplankton assembling

In the lowest size range, each dataset displays an undersampling (Fig S7a) which is symptomatic of either incorrect detection of objects due to optical or digital limitation of each device (e.g. ^72^) or, when using nets, to mesh extrusion of organisms. Therefore, any parts of the NBSS and BSS below the maximal abundance of each device were discarded before assembling them. For the highest size range of each dataset, very large organisms correspond to a presence-absence signal rather than quantitative due to insufficient sampling effort, and were disregarded. Symptoms of such observations are recurrent size bins with observations corresponding to 1-2 organisms surrounded by multiple empty bins. For this we take the objective criteria that every NBSS size bin separated by more than 5 empty size bins were disregarded.

Three different ways of merging all observations were considered. In all cases we considered the principle that, when represented in logarithmic scale, the intercept of NBSS spectra represents the total abundance of organisms in the considered ecosystem^73,74^. Therefore, discrepancies in intercepts only reflect discrepancies in sampling such as different depths or strategies (discrete vs integrative).

Since WP2 net observations were among the more commonly sampling devices used throughout the campaign (see Fig. S1), but also cover an intermediate size within observations, we used them as a global reference. Therefore, without WP2-net observations, no adjustment was performed, and data were not considered in the analysis. Only NBSS of total living organisms were considered.

1) intercept-adjustment

The first, and preferred correction method directly relies on the theory: using log transformed biovolume and NBSS data, we estimated the intercept and slope on WP2 net observations (WP2_i_ and WP2_s_ respectively). The intercept on other datasets (Dataset_i_) was calculated by imposing those measured in the WP2 (Fig S7c) and the NBSS of each dataset is corrected by a factor which corresponds to the intercept discrepancy observed such as: NBSS_corr_= NBSS_raw_*exp(WP2_i_-Dataset_i_)

Such correction was in most cases sufficient to effectively correct for intercept discrepancies.

However, it was inadequate in specific cases such as when multiples bumps were observed either on the NBSS from the WP2 net or on other datasets, therefore compromising slopes or intercepts estimates, or when the dataset considered does not span large size ranges (mostly from UVP or IFCB observations).

2) Default adjustment

Secondly, given the overlap in size between instruments, some overlap observations could be present. An adjustment ratio is computed for each overlapping NBSS size bins from which a median conversion ratio between each pair of size-overlapping instruments could be calculated. Each of those ratios corresponds roughly to the intercept correction as mentioned above. All these adjustments between instruments were accumulated across stations to produce a median ratio of correction which was applied to sequentially correct each series of observations to a comparable level with WP2 observations (Fig S7b).

3) Site-specific adjustment

Finally, a site-specific adjustment was produced, using, if present, the median correction ratio specifically observed at that given site or the default ratio if no specific correction was present (Fig S7d). This one was usually preferred to the default adjustment.

#### Final adjustment

Results for all adjustments were inspected to detect any slope breaks in the final NBSS (Fig S7f) and BSS spectra (Fig S7i), these latter being symptomatic of incorrect corrections. If the intercept-adjustment was qualified as inadequate, other possibilities were tested to obtain the best final adjustment (Fig S7e). In total, for day observations, we obtained metaplankton assemblages from 11 different dataset sources, which corresponded to 695 datasets adjusted with 152 WP2 net observations. The intercept adjustment was adequate for 529 datasets (76.1%) while site-specific and default adjustments were applied on 99 (14.2%) and 52 (7.48%) datasets respectively. For e-HFCM datasets for which neither default nor specific adjustments were possible (no shared size classes with WP2 nets), the datasets were kept un-corrected, and this occurred in for 15 cases (2.15%).

Final corrections were applied to obtain NBSS and BSS for the total living organisms, but also for combining observations done at the functional and trophic levels (Fig S7g-l). For these ones, at a given size bin, a mean between the different datasets was performed. No assemblages were performed at the initial taxonomic identification level because the variations of taxonomic level of identification between datasets may lead to duplicate counts (e.g. “copepods” identified with UVP correspond to several families of copepods identified with Zooscan). Finally, biovolumes and numbers of organisms were summed across sizes to provide an overview of the contribution of the different functional groups and trophic levels to the full metaplankton assemblage.

The availability of the various data sources (Fig S1) varies across the expedition. Hence, metaplanktonic assemblages were obtained with different granularities:

1. “***Meta-Plk >0.8 µm***” spans organisms ranging from 0.8 µm to several cm (including flow-cytometry, IFCB, Flowcam, Zooscan from several nets and UVP and with a complete coverage in between), it is only available from the Arctic ecosystem and covers 20 sites.
2. “***Meta-Plk >20 µm***” spans organisms ranging from 20 µm to several cm (using e-HFCM with 20 µm net, Zooscan from WP2 and Regent nets and UVP). It covers both polar and tropical parts of the expedition and includes 63 stations.

Two other more heterogeneous products were generated for a more complete global geographic coverage, although these could suffer from higher uncertainties due to their incomplete coverage: “***Meta- Plk >0.8 µm incomplete***” is similar to the above but miss some observation for particular organism sizes, notably due to the absence of Flowcam analyses of the 20 µm net fractions. “***Meta-Plk heterogeneous***” regroups all observations available at a given site with heterogeneous coverage in size classes but when compared with more complete datasets (Fig S2, S3), provides an independent confirmation over wider geographic coverage.

### Carbon biomass calculations

Biovolume estimates were converted to carbon biomass by using conversion factors from several sources. For most phytoplankton and microzooplankton, we combined conversion factors between biovolume and carbon biomass^75–84^ to obtain a single usable relationship to convert biovolume estimates to carbon biomass (Table S5, Fig. S8) following the relationship:

Biomass (mgC)= a * Biovolume (mm3) ^b.

For rhizarians including acanthareans, foraminifers, phaeodaria and radiolarians, we used conversions between biovolume and carbon biomass^44,85^ except for colonial and solitary collodarians, Spumelarians, nasselarians and specific genus of phaeodarians (Alaucantha and Protocystis) for which a specific carbon to biovolume relationships were used instead^86^. Finally for larger zooplankton we used phylum specific conversion factors between wet mass and carbon mass^87^ assuming that biovolume estimates are comparable to wet mass. All conversion factors for each taxonomic identification are presented in Table S4. This conversion work allowed us to check if our observations, mostly expressed in biovolume, are robust even when expressed in carbon (Fig 2, 4, S3, S4). We are however conscious that such conversion may also introduce biases. It is worth noting that while carbon units are widely representative of the respiration expenditures of organisms, their wet mass (and biovolume) is a better reflection of their feeding activity and interactions even when considering gelatinous plankton^49^, and therefore biovolume is preferred here in the context of trophic structure.

### BSS, NBSS and trophic slopes calculations

On metaplanktonic data we calculated three different slopes characterising the ecosystem structure (Fig. 1). Both BSS and NBSS spectral slopes (*SBSS* and *SNBSS*) were calculated on biovolume (mm^3^ m^-3^ and mm^3^ mm^-3^ m^-3^ respectively; Fig. 1 S7) as a function of the median biovolume (mm^3^) of each size class.

Similar calculations were done in carbon units (mgC mgC^-1^ m^-3^) as a function of the median carbon mass (mgC) of each size class. All data were log transformed and linear adjustments were obtained on log transformed data. For the trophic slope (*STrophic*) calculations, total biovolume (mm^3^ m^-3^) or carbon mass (mgC m^-3^) of each trophic group were summed for each station and attributed respectively to trophic levels 1, 1.5, 2, 2.5 and 3 (Fig. 1, Fig. S7). Total biovolumes or biomasses were log-transformed and the trophic slope was calculated as the slope of these log-transformed biovolume/biomass as a function of trophic level.

For any linear and power relationships used in the manuscript, we used robust linear fitting which is less sensitive to possible outliers^88^.

To further analyse the trophic structures of plankton communities worldwide, we classified bottom-heavy, flat, and top-heavy food webs as having trophic slopes STrophic ∈ (< -0.25, -0.25 to 0.25 and > 0.25) or, alternatively, SNBSS ∈ (<-1,1 ; -1.1 to -0.9 ; > -0.9).

### Genomic data

DNA metabarcoding data, which target the Eukaryota kingdom through the V9 region of the 18S rRNA gene, were used in this study to assess if the trends observed for eukaryotes with imaging approaches were consistent with those based on molecular data. A full description of all the steps from sampling to bioinformatic analysis leading to the OTU table are described in^2,7^ and available in^89,90^. The number of reads associated with each OTU was used as a proxy of abundance for our ecological analysis. The number of reads in a sample does not reflect total biomass/abundance variations, we therefore standardized the read counts dividing by the total number of reads in each sample. Mesoplanktonic subsurface samples (180-2000 µm; the biggest size fraction for metabarcoding in *Tara* Oceans) from surface samples were selected for the comparison (136 samples) for their good size overlap with nets used for imaging samples. We assigned these OTUs to functional and trophic groups compatible with the one used for imaging datasets. The complete association with the different functional and trophic status of organisms can be found in Table S6. Additional trophic groups (bacteriophages, parasitic) were also considered and relative read abundance for each functional and trophic group was calculated at the sample level but they were not included in the trophic slope calculation. We calculated a trophic slope over the trophic groups 1-3 (autotrophs to carnivores) by first log-transforming the relative reads and by calculating the slope of the log-transformed relative counts as a function of trophic level (Fig. S4e). It should be noted that the interpretation of metabarcoding-derived slopes is subject to caution due to PCR biases. Indeed, the correlation of this trophic slope with quantitative imaging methods is not significant (Fig S1), but it gives relatively similar proportion of top-heavy ecosystems worldwide, with comparable functional composition (Fig 2a), which both follow the same geographical pattern as imaging observations (Fig 2b, c, S4).

### Environmental data

To interpret our observations, we used combined environmental data representative from each station. All our observations span several samples done during the 1-2 days of sampling of the station. The net tows and the UVP casts were vertically integrated. We therefore compiled data relying on published datasets^56^ that correspond to water column features^91^ and mesoscale features^92^ that we enriched with data corresponding to nutrients levels^93^, carbonate chemistry^94^ and pigments concentrations^95^. All these data correspond to the median of sample values. To combine information relevant to the entire site and water column, we did restrict the dataset to samples encompassing at least 0-50m depth integration and generated a single mean for the site. We further enrich this contextual data by calculating carbon fluxes obtained from the UVP data^96^ and averaging around 2 specific depths (200 and 500m ±20m). From pigment composition, we derived chlorophyll *a* into micro, pico and nanoplankton proportions (micro, pico and nano) using the Uitz et al^97^ algorithm. We also calculated and index of nitrate deficiency relatively to phosphate (N*) using the following formula^98^:

N*=Nitrate + Nitrite – (16 * Phosphate)

The environmental parameters ultimately used in our analysis include concentrations and limitations by different nutrients (N, P, Si, Fe, N*), the amount of photosynthetically available radiations (PAR), some water column information such as the mixed layer depth (Zmld) and the euphotic depth (Zeu), temperature and salinity of the euphotic zone (Tzeu and Szeu), mesoscale indices such as the Okubo– Weiss parameter (okubo) that indicates if the station is located within an eddy (negative value) or outside an eddy (positive value), the Lyapunov exponent correlated with the stability of movements at the mesoscale level and the residence time which indicates how many days a water mass has spent inside an eddy. Finally, a certain number of parameters relative to the biological context and productivity of the ecosystems were also used: the sea-surface chlorophyll a concentration at 10m (chla ss), the net primary production (NPP), carbon flux at 200 and 500m (flux200, flux500) and the percentage of chlorophyll a represented by micro-, pico- and nano-plankton (micro, pico, nano).

### Statistical analysis

To interpret how stations are characterized by their plankton functional types composition, we performed a principal component analysis (PCA) using the functional groups composition (using Hellinger transformation). Groups of stations sharing similar compositions were established on the two first axes of the PCA, using Euclidean distances (i.e. Hellinger distances) and Ward linkage. The association with the environment was tested by adding environmental variables as supplementary variables in the analysis and evaluating their correlations to the PCA components.

## Acknowledgements

*Tara* Oceans (which includes both the *Tara* Oceans and *Tara* Oceans Polar Circle expeditions) would not exist without the leadership of the Tara Ocean Foundation and the continuous support of 23 institutes (http://oceans.taraexpeditions.org). We further thank the commitment of the following sponsors: CNRS (in particular Groupement de Recherche GDR3280 and the Research Federation for the study of Global Ocean Systems Ecology and Evolution, FR2022/Tara Oceans-GOSEE), European Molecular Biology Laboratory (EMBL), Genoscope/CEA, The French Ministry of Research, and the French Government ‘Investissements d’Avenir’ programmes OCEANOMICS (ANR-11-BTBR-0008), FRANCE GENOMIQUE (ANR-10-INBS-09-08), MEMO LIFE (ANR-10-LABX-54), and PSL* Research University (ANR-11-IDEX-0001-02). Funding for the collection and processing of *Tara* Oceans data set was provided by NASA Ocean Biology and Biogeochemistry program under grants NNX11AQ14G, NNX09AU43G, NNX13AE58G and NNX15AC08G to the University of Maine, and Canada Excellence Research Chair on Remote sensing of Canada’s new Arctic frontier and Canada Foundation for Innovation. We also thank the support and commitment of Agnès b. and Etienne Bourgois, the Prince Albert II de Monaco Foundation, the Veolia Foundation, Region Bretagne, Lorient Agglomeration, Serge Ferrari, Worldcourier, and KAUST. The global sampling effort was enabled by countless scientists and crew who sampled aboard the *Tara* from 2009-2013, and we thank MERCATOR-CORIOLIS and ACRST for providing daily satellite data during the expeditions. We are also grateful to the countries who graciously granted sampling permissions. We thank the EMBRC platform PIQv for image analysis. This work was supported by EMBRC-France, whose French state funds are managed by the ANR within the Investments of the Future program under reference ANR-10-INBS-02. FL thanks the Institut Universitaire de France (IUF) and support from NOAA Award NA21OAR4310254. FL and LG had received funding from the European Union’s Horizon 2020 research and innovation programme “Atlantic Ecosystems Assessment, Forecasting and Sustainability” (AtlantECO) Grant ID: 862923. FL and CdV had received co-funding by the European Union (GA#101059915 - BIOcean5D). ICM-CSIC authors were supported by the Severo Ochoa Centre of Excellence’ accreditation (CEX2019-000928-S). CB acknowledges funding from the European Research Council under the European Union’s Horizon 2020 research and innovation programme (grant agreement 835067 (Diatomic)). MCB acknowledges funding from the Coordination for the Improvement of Higher Education Personnel of Brazil (CAPES 99999.000487/2016-03) and the French Facility for Global Environment (FFEM). Support from NASA’s Ocean Biology and Biogeochemistry program. ML was supported by the TULIP Laboratory of Excellence (ANR-10-LABX-41). We also thank the many different students that helped to taxonomically annotate the image which includes Llopis-Monferrer Natalia, Olivier Marion, Etienne Dvorak, and Madeline Carsique.

Views and opinions expressed are however those of the author(s) only and do not necessarily reflect those of the European Union. Neither the European Union nor the granting authority can be held responsible for them. The authors declare that all data reported herein are fully and freely available from the date of publication, with no restrictions, and that all of the analyses, publications, and ownership of data are free from legal entanglement or restriction by the various nations whose waters the *Tara* Oceans expeditions sampled in. This article is contribution number XX of *Tara* Oceans.

## Authors contribution

FL designed the study, taxonomic annotation, data analysis, and wrote the paper, LG collected *Tara* Oceans samples, analysed oceanographic data, designed the study, participated to data analysis and to the manuscript writing, MCB taxonomic annotation of the regent dataset; provided constructive comments, revised and edited the manuscript, LPC generated and provided the e-HFCM >20µm dataset, SC generated the e-HFCM >5µm dataset, constructed, provided taxonomic annotation of the e-HFCM >5µm dataset, JRD analysed taxonomic data to assign trophic levels to taxa, AL constructed, provided and curation of the WP2, bongo and regent datasets, JMG processed all the Facscalibur flow cytometry, curation and quality control, processed the data to extract size provided constructive comments, revised and edited the manuscript, PLG constructed, provided and taxonomic annotation of the IFCB dataset, NH constructed, provided and annotated the functions and trophic levels in the meta-B datasets, FMI provided taxonomic annotation of the e-HFCM >20µm dataset, provided constructive comments, revised and edited the manuscript, LJ taxonomic annotation of the WP2, bongo and regent datasets, quality control of metadata and taxonomic identifications, ML provided constructive comments, revised and edited the manuscript, SM provided constructive comments, revised and edited the manuscript, ZM provided statistical analysis, graphical help, revised and edited the manuscript, MP designed the sample collection, collected *Tara* Oceans samples, analysed oceanographic data, ensured quality control on imaging datasets, helped in the import/export from EcoTaxa from various instruments, JJPK annotated the e-HFCM >20µm dataset, RP designed the microscopic frame to generated the e-HFCM >5µm dataset, provided funding, J-BR constructed, provided, curation and taxonomic annotation of the WP2, bongo and regent datasets, LZ provided constructive comments, revised and edited the manuscript, LS collected *Tara* Oceans samples, designed the sampling and provided funding and supervision of the ZooScan and UVP datasets, SA collected samples, provided the Facscalibur flow cytometry, LK-B collected *Tara* Oceans samples, analysed oceanographic data; provided constructive comments, supervised the analysis of IFCB and Flowcam datasets, revised and edited the manuscript, EB collected *Tara* Oceans samples, analysed oceanographic data, provided constructive comments, revised and edited the manuscript, MBS provided constructive comments, revised and edited the manuscript, CdV collected samples, generated and provided the e-HFCM >5µm and DNA metaB datasets, provided constructive comments, revised and edited the manuscript, supervised the study, CB collected samples, provided constructive comments, revised and edited the manuscript, supervised the study, EK collected samples, supervised the study, GG collected samples, provided constructive comments, revised and edited the manuscript; supervised the study. Tara Oceans coordinators provided constructive criticism throughout the study.

Consortium

Tara Oceans Coordinators and Affiliations

Silvia G. Acinas8, Marcel Babin21, Peer Bork6,22,23, Emmanuel Boss17, Chris Bowler2,11, Guy Cochrane24, Colomban de Vargas2,19, Michael Follows25, Gabriel Gorsky1,2, Nigel Grimsley2,26,27, Lionel Guidi1,2, Pascal Hingamp2,28, Daniele Iudicone29, Olivier Jaillon2,30, Stefanie KandelsLewis3,20, Lee Karp- Boss5, Eric Karsenti2,6,16,29, Fabrice Not2,19, Hiroyuki Ogata31, Stéphane Pesant32, Nicole Poulton33, Jeroen Raes34,35,36, Christian Sardet1,2, Sabrina Speich 2,37,38, Lars Stemmann1,2, Matthew B. Sullivan18, Shinichi Sunagawa39, Patrick Wincker2,30

21Département de biologie, Québec Océan and Takuvik Joint International Laboratory (UMI 3376), Université Laval (Canada) - CNRS (France), Université Laval, Québec, QC, G1V 0A6, Canada

22Max-Delbrück-Centre for Molecular Medicine, 13092 Berlin, Germany

23Department of Bioinformatics, Biocenter, University of Würzburg, 97074 Würzburg, Germany

24European Molecular Biology Laboratory, European Bioinformatics Institute (EMBL-EBI), Welcome Trust Genome Campus, Hinxton, Cambridge, UK

25Department of Earth, Atmospheric, and Planetary Sciences, Massachusetts Institute of Technology, Cambridge, MA 02139, USA

26CNRS UMR 7232, Biologie Intégrative des Organismes Marins, Avenue du Fontaulé, 66650 Banyuls-sur-Mer, France

27Sorbonne Université Paris 06, OOB UPMC, Avenue du Fontaulé, 66650 Banyuls-sur-Mer, France

28Aix Marseille Univ., Université de Toulon, CNRS, IRD, MIO UM 110, 13288, Marseille, France

29Stazione Zoologica Anton Dohrn, Villa Comunale, 80121 Naples, Italy

30Génomique Métabolique, Genoscope, Institut François Jacob, CEA, CNRS, Univ Evry, Université Paris-Saclay, 91057 Evry, France

31Institute for Chemical Research, Kyoto University, Gokasho, Uji, Kyoto 611-0011, Japan

32EMBL’s European Bioinformatics Institute (EMBL-EBI), Hinxton, UK

33Bigelow Laboratory for Ocean Sciences, East Boothbay, ME, 04544, USA

34Department of Microbiology and Immunology, Rega Institute, KU Leuven, Herestraat 49, 3000 Leuven, Belgium

35Center for the Biology of Disease, VIB KU Leuven, Herestraat 49, 3000 Leuven, Belgium 36Department of Applied Biological Sciences, Vrije Universiteit Brussel, Pleinlaan 2, 1050 Brussels, Belgium

37Department of Geosciences, Laboratoire de Météorologie Dynamique (LMD), Ecole Normale Supérieure, 24 rue Lhomond 75231 Paris, Cedex 05, France

38Ocean Physics Laboratory, University of Western Brittany, 6 avenue Victor-Le-Gorgeu, BP 809, Brest 29285, France

39Institute of Microbiology, ETH Zurich, Zurich, Switzerland

## Data availability

*Imaging:* EcoTaxa (see table SI-3) + ZENODO repository for tsv used + post-processed data + intercalibrated assembled measurements

https://zenodo.org/records/10478781

*Metabarcoding data:*

samples= https://zenodo.org/record/3768510#.XtjE9Z4zb1I references https://zenodo.org/record/3768951#.XtjUdJ4zb1I *Environmental data*: Pangea (see methods)

## Supplementary information

This part describes in detail the specificity of each optical or imaging equipment that was used during the *Tara* Oceans sampling.

### Accuri

Starting from station 154 (i.e. temperate to polar stations), samples used for flow cytometry were collected with Niskin bottles (0-3m). Samples were analysed alive on-board using an Accuri flow cytometer (BD Accuri C6). Each sample was run twice, in fast mode (163.5µl / min; sample size of 327µl) and in slow mode (33µl / min; sample size of 66µl) for optimal detection and counts of large and small particles. Calibration of fluorescence peaks (BD, 8 & 6 µm validation beads) and counts (1µm Polyscience yellow beads) were done daily. Size calibrations were done weekly using calibration beads (1, 2, 4, and 10µm) but also using 13 phytoplankton cultures of known sizes, ranging from 1-25 µm from which we created a calibration curve to estimate cell sizes of natural phytoplankton populations (Size (µm)=FSC*0.0000041+0.85). Size of cultures >3 µm were confirmed by measuring cell size under the microscope. Blanks of filtered seawater samples were run with each set of samples and the background signal was gated and removed from each sample to ensure that only populations of cells were counted. Flow cytometry data were gated and for the sake of simplicity we chose to only gate out all particles that were considered as not alive without separating the different populations.

### FACSalibur flow cytometry

Sampling and analysis are described in^1^. We restricted our analysis to sea-surface samples collected using Niskin bottles. Available counts are variable within stations notably the the nano-eukaryotes counts added in the Arctic part of the expedition (Fig S1). Concentration and mean size of the different cell populations detected were measured and used to derive an equivalent spherical biovolume. Since the individual size of each cell was not available, the size range (minimal-maximal size) of each population was not available, it was then not possible to size-normalize those counts and therefore could not be integrated in the NBSS approach. However, since NBSS results correspond roughly to concentrations^2^, we converted raw concentrations by the scaling factor Bv_mean_/Bv_range_ to obtain comparable units with NBSS spectra for gross cross comparison display (Fig S7), but we did not use them to calculate NBSS slopes. The total biovolume observed by flow cytometry of each category was also summed to provide an overview of the full (0.2-cm size) composition and trophic structure of plankton. This estimation was only done when both bacterial and photosynthetic picoplankton were available. Since this approach cannot be homogenized with other measurements, a certain bias could have been introduced, however it confirmed that extending the range of observation from 20 µm down to 0.2 µm did not change most of our observations and findings (Fig S5, S6b)

### eHFCM H5/H20

The environmental High Content Fluorescence Microscopy^3^ (e-HCFM) is a 3D multichannel imaging workflow which was applied on samples originating from the 5 and 20µm nets. Protocols for acquisition were described in^3^. Briefly, it allows to take confocal images at various focal distance (Z-stacks) using 5 different excitation channels (Bright field, and 4 fluorescence channels looking for specific stainings such as Hoechst33342-DNA staining; Poly-L-lysine-Alexa Fluor staining for external membranes, proteins and structures, DiOC6(3) staining internal membranes and for chlorophyll autofluorescence). For all objects, single layer images were constructed and all morphological measurements together with associated metadata were imported to EcoTaxa. As with other instruments, we used the major and minor axis of every image to calculate their ellipsoidal equivalent biovolume. Since a 5µm or 20µm net was used for each of those datasets, we disregarded every particle below 3 and 12µm, respectively, which often corresponds to artefacts or fragments generated during the preparation process.

### IFCB

The Imaging FlowCytoBot (IFCB^4,5^) was connected to the inline system and imaged approximately a 5mL sample of seawater every 25 minutes. The IFCB was set to record images for all particles above a Chl *a* in vivo fluorescence trigger level, therefore ignoring other particles. All images were saved together with various measurements by the instrument itself. All images were processed with a publicly available custom MatLab code (https://github.com/hsosik/ifcb-analysis) and exported together with associated metadata to EcoTaxa^6^ for taxonomic identification (https://github.com/OceanOptics/ifcb-tools). We directly used the “summed biovolume” calculated by the IFCB to extract the biovolume of each organism.

### FlowCam

Samples from Niskin bottles and from the 20µm net were analyzed on-board using the FlowCam analyzer^7^ (Fluid Imaging Technologies; model Benchtop B2 Series equipped with a 4X lens). The FlowCam is an automated microscope taking images while organisms are pumped through a capillary imaging chamber. Here we used the auto-trigger mode to image the particles in the focal plane at a constant rate. Raw images were analyzed using Zooprocess software (https://sites.google.com/view/piqv/zooprocess) which allows to subtract the background, detect and measure different morphological characteristics of imaged particles and store the vignettes of every detected object > 20µm. All the images and associated metadata were imported to EcoTaxa for taxonomic identification. We used the major and minor axis of every imaged object to calculate its ellipsoidal equivalent biovolume.

### Nets -Zooscan

All samples originating from nets >200µm were fixed on-board with borax-buffered formaldehyde (3.5% final volume) and analyzed on land. For the analysis, the sample was gently filtered (100µm mesh) and transferred to filtered seawater. WP2 and bongo net samples were separated into two size classes 100-1000µm and >1000µm and only a single fraction was considered for the Regent net.

Fractions were split using a Motoda box^8^ and a subsample containing approximately 1000 objects was scanned using a Zooscan system^9^. This sampling strategy allows to correctly take into account both the numerous small organisms and the rare large ones. The scans were processed using the Zooprocess software. All images and associated metadata were imported to EcoTaxa for taxonomic identification. We used the major and minor axis of every image to calculate their ellipsoidal equivalent biovolume.

### UVP

The Underwater Vision Profiler^10^ (UVP-5) is an underwater imager mounted on the RVSS. This system allows to illuminate a precisely calibrated volume of water and capture images at a rate of 20 images s^-1^ during the descent. The recorded images were treated via the Zooprocess software as described above and particles >100µm were detected, counted and measured and were considered as marine snow. Particles >600µm were imported to EcoTaxa as vignettes with associated metadata and sorted for taxonomic classification. As done with other instruments, we used the major and minor axis of every image to calculate their ellipsoidal equivalent biovolume.

## Supplementary information

**Figure S1:**
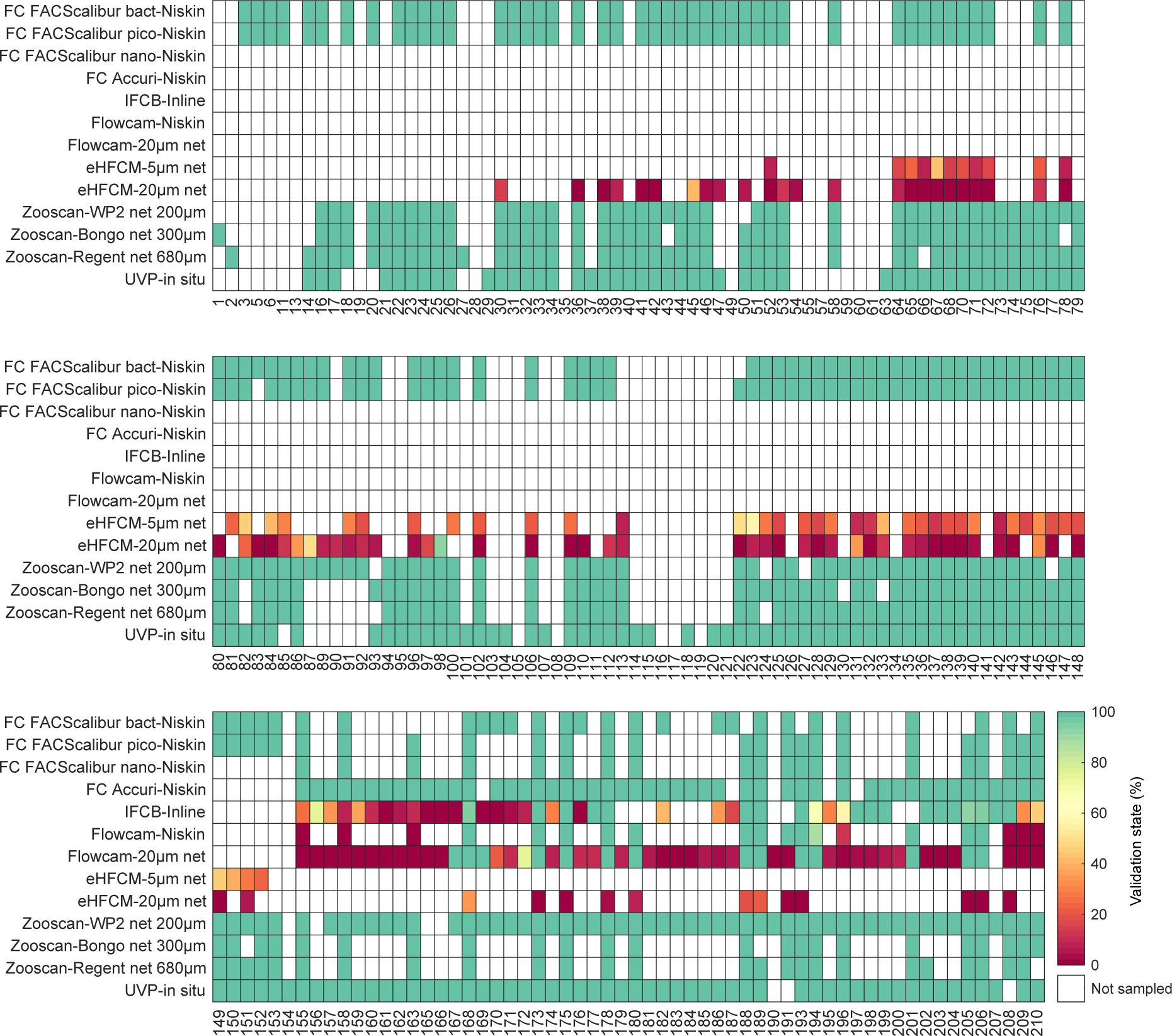
Availability and taxonomic validation state (percentage of predicted taxonomical annotation validated by taxonomic expert) of the imaging datasets across the 210 sampling stations. Note the increased availability of instruments for the TOPC Arctic expedition.

**Figure S2:**
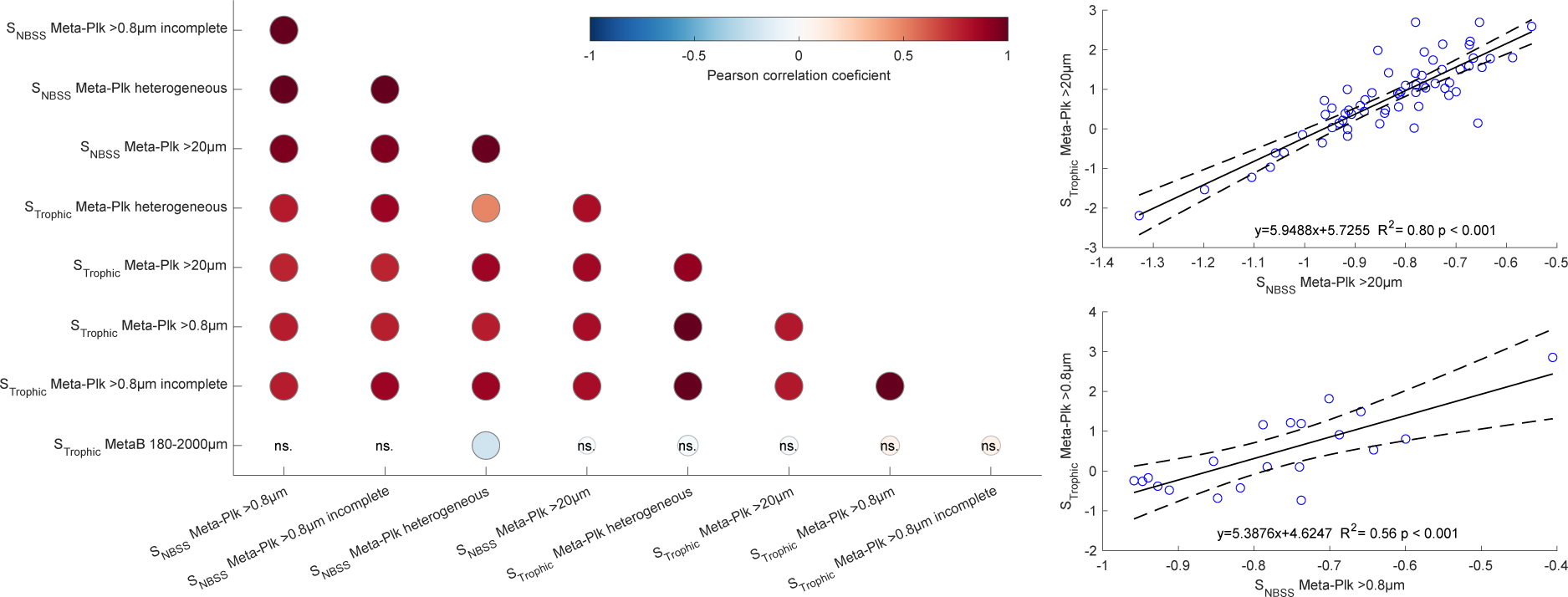
Correlations between slopes indicating the structure of planktonic ecosystems. a) Spearman’s rank correlation coefficients between NB-SS and Trophic slopes calculated from the different levels of aggregation of metaplanktonic assemblages. b) Correlation between the NB-SS and Trophic slopes calculated from the metaplanktonic assemblage >0.8µm (mostly polar) and b) >20µm (mostly tropical).

**Figure S3:**
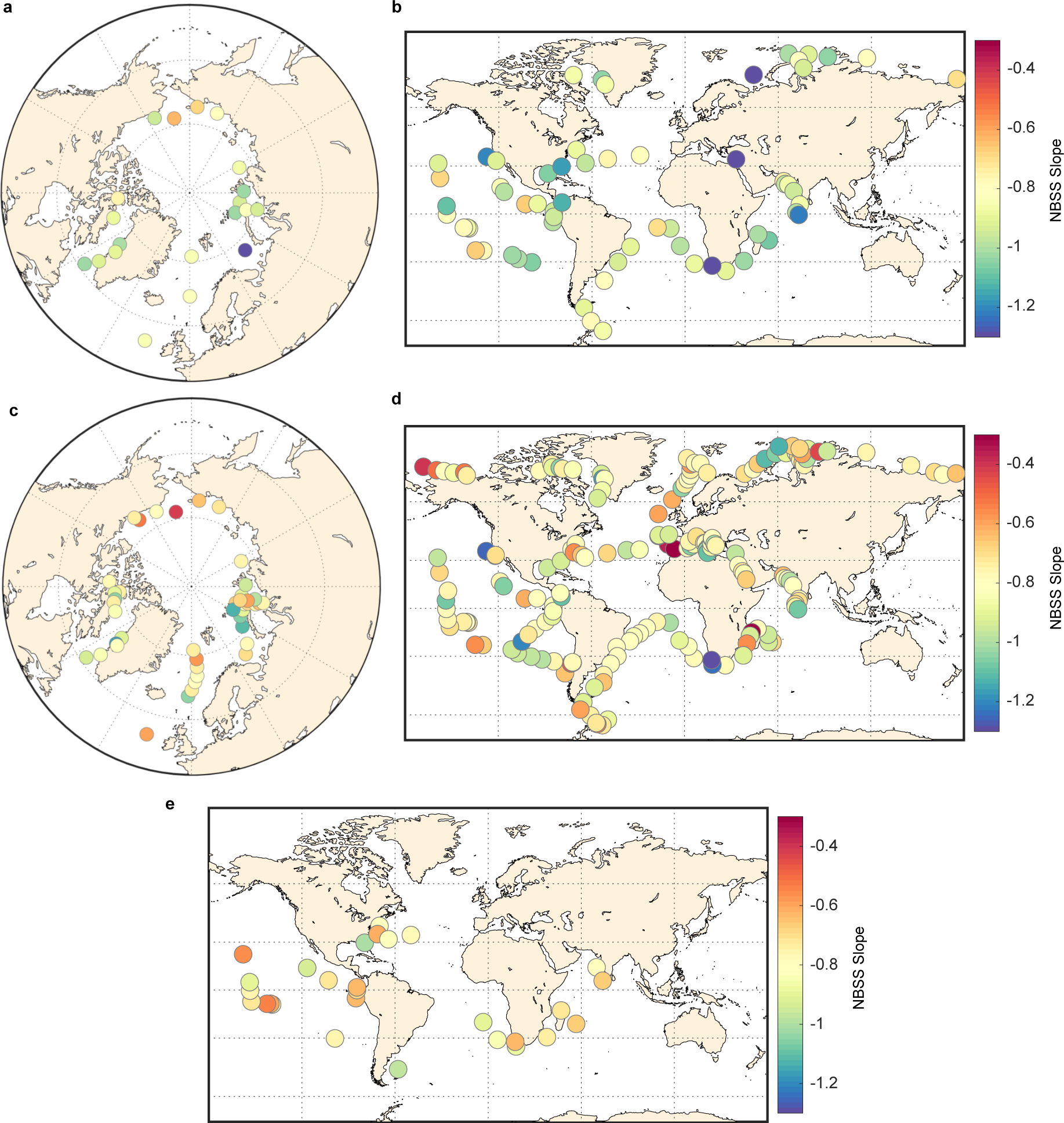
Slopes of the normalized biomass size spectra (NB-SS) using carbon weight for a) the Arctic ecosystem obtained with Meta-Plk >0.8 µm or b) for the Meta-Plk >20µm at the global scale. In both cases a large majority of observations have slopes corresponding to top-heavy trophic structures. NB-SS using biovolumes for incomplete and heterogeneous versions of the datasets c) Meta-Plk >0.8 µm incomplete or d) Meta-Plk heterogeneous or e) Meta- Plk >20 µm at night-time.

**Figure S4:**
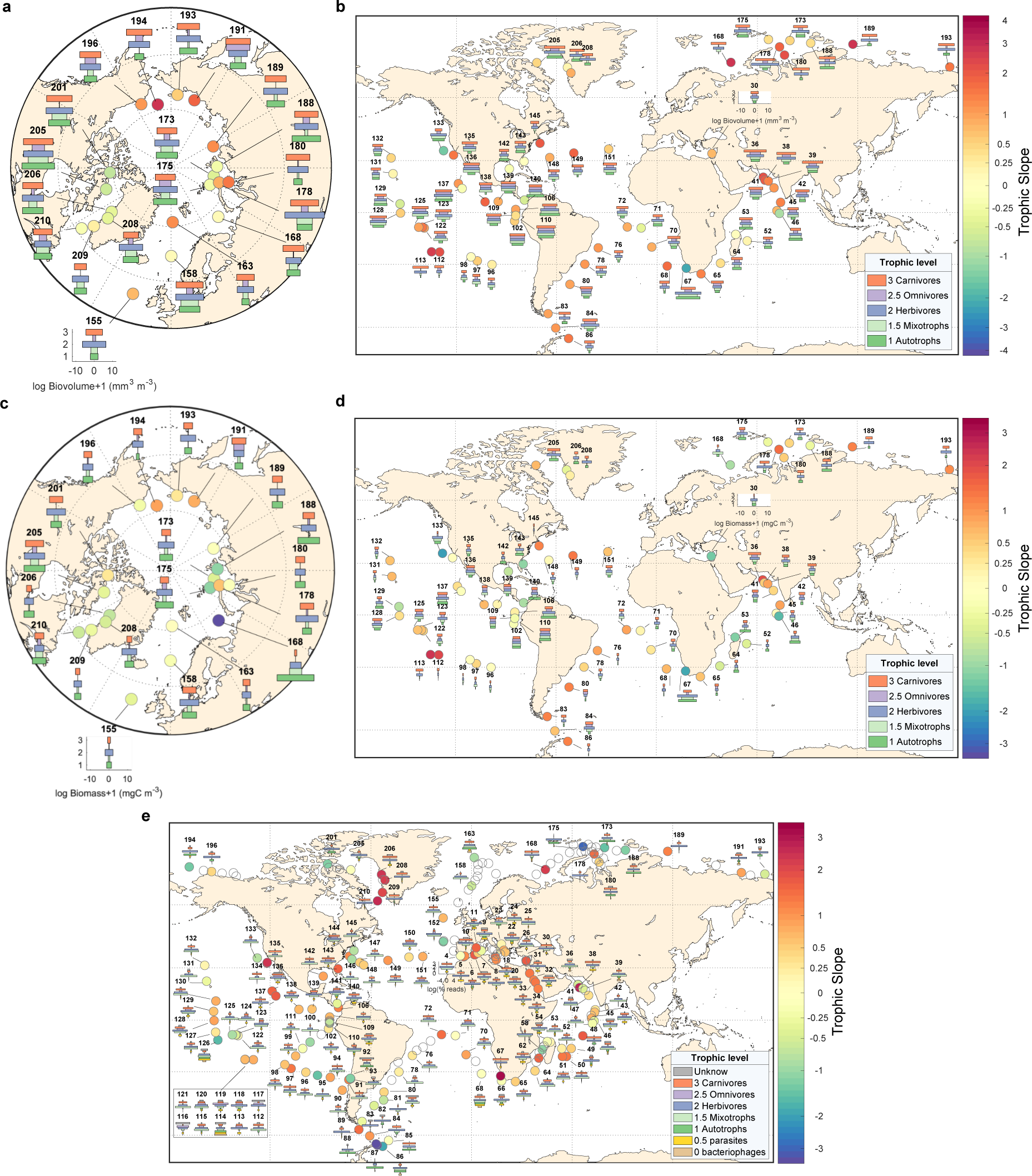
Trophic pyramids and trophic slopes obtained from the different metaplankton producsa) Total biovolume of the Meta-Plk > 0.8µm assemblage split between the different trophic levels and resulting trophic slope. b) Same as a) but with Meta-Plk > 20 µm assemblage. c) and d) Same as a & b but expressed in carbon units. e) Same as a) but using total reads of meta-barcoding originating from surface nets from size fraction 180- 2000µm.

**Figure S5:**
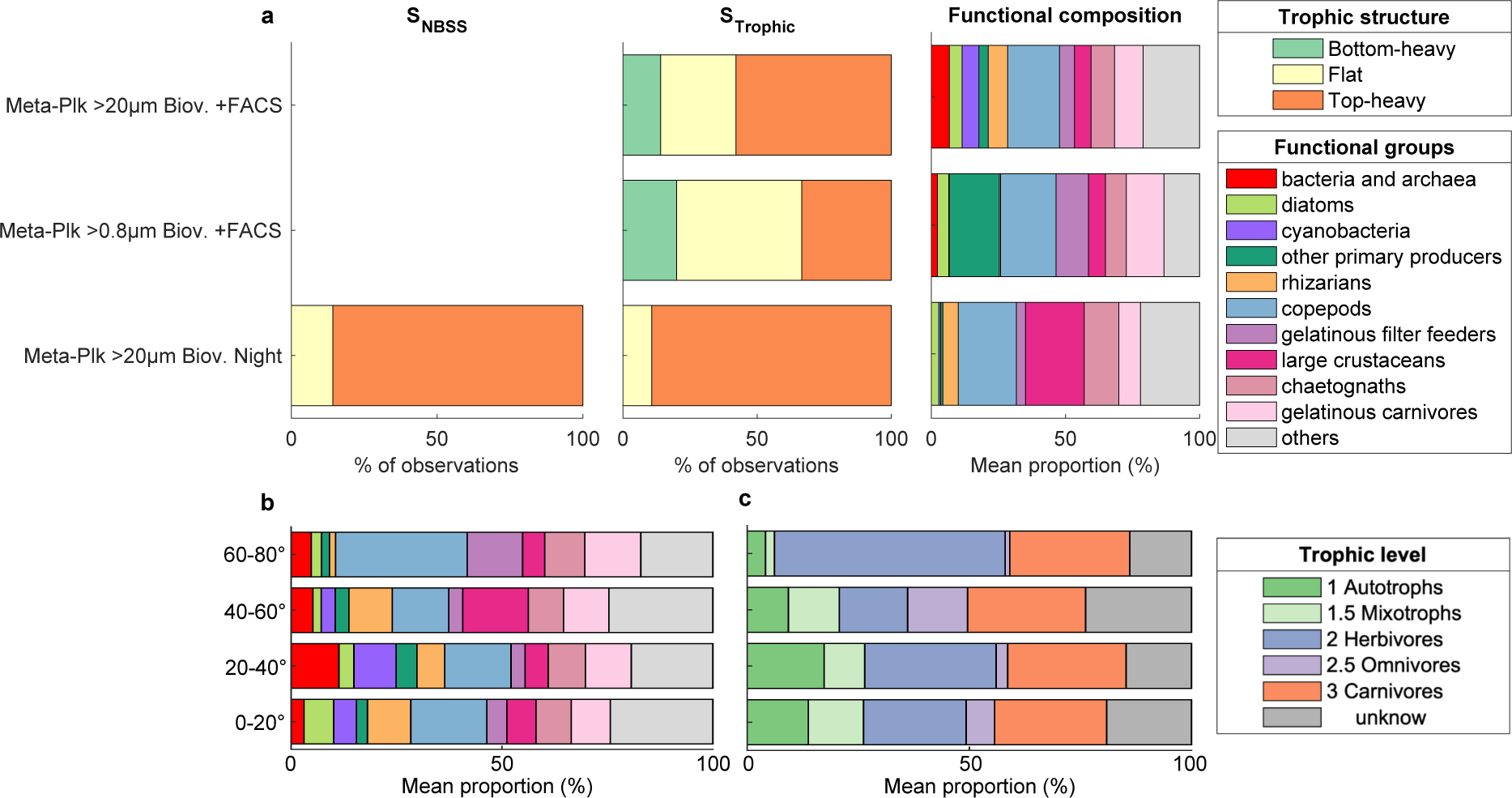
Same as Figure 2 (proportions of biovolume) but testing alternative results with adding FACScalibur flow cytometry observations to either global >20µm Metaplankton reconstruction or to the arctic >0.8µm Metaplankton or testing results obtained at night time. Latitudinal variations of the different b) functional groups or c) trophic groups of plankton obtained with adding the FACScalibur flow cytometry observations to global >20µm Meta-plankton reconstruction.

**Figure S6:**
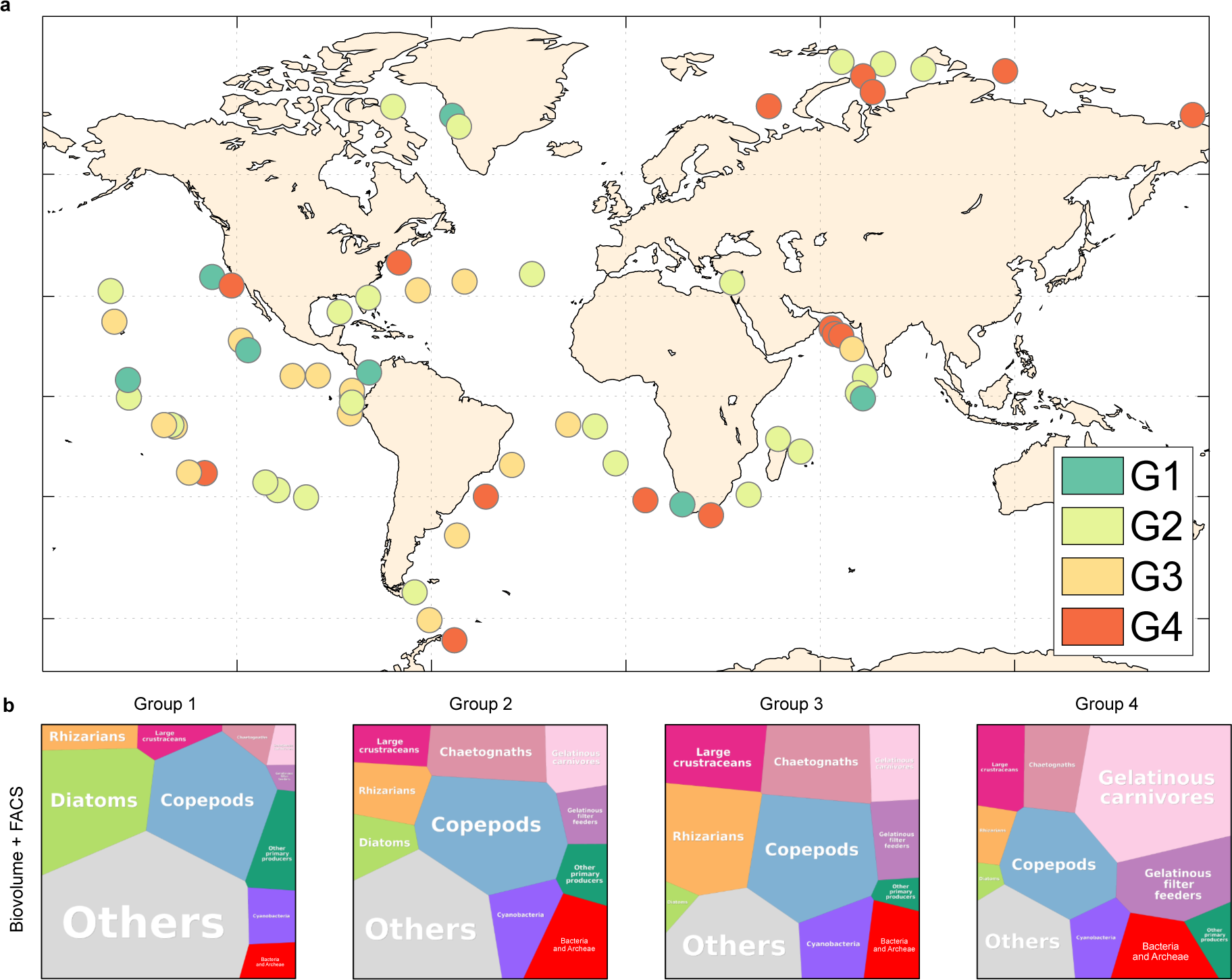
a) Position of stations identified as belonging to the different groups of stations based on their functional composition (Fig. 4). b) Mean biovolume composition of these groups of stations when adding FACScalibur Flow Cytometry observations in the global >20µm Meta-plankton reconstruction.

**Figure S7:**
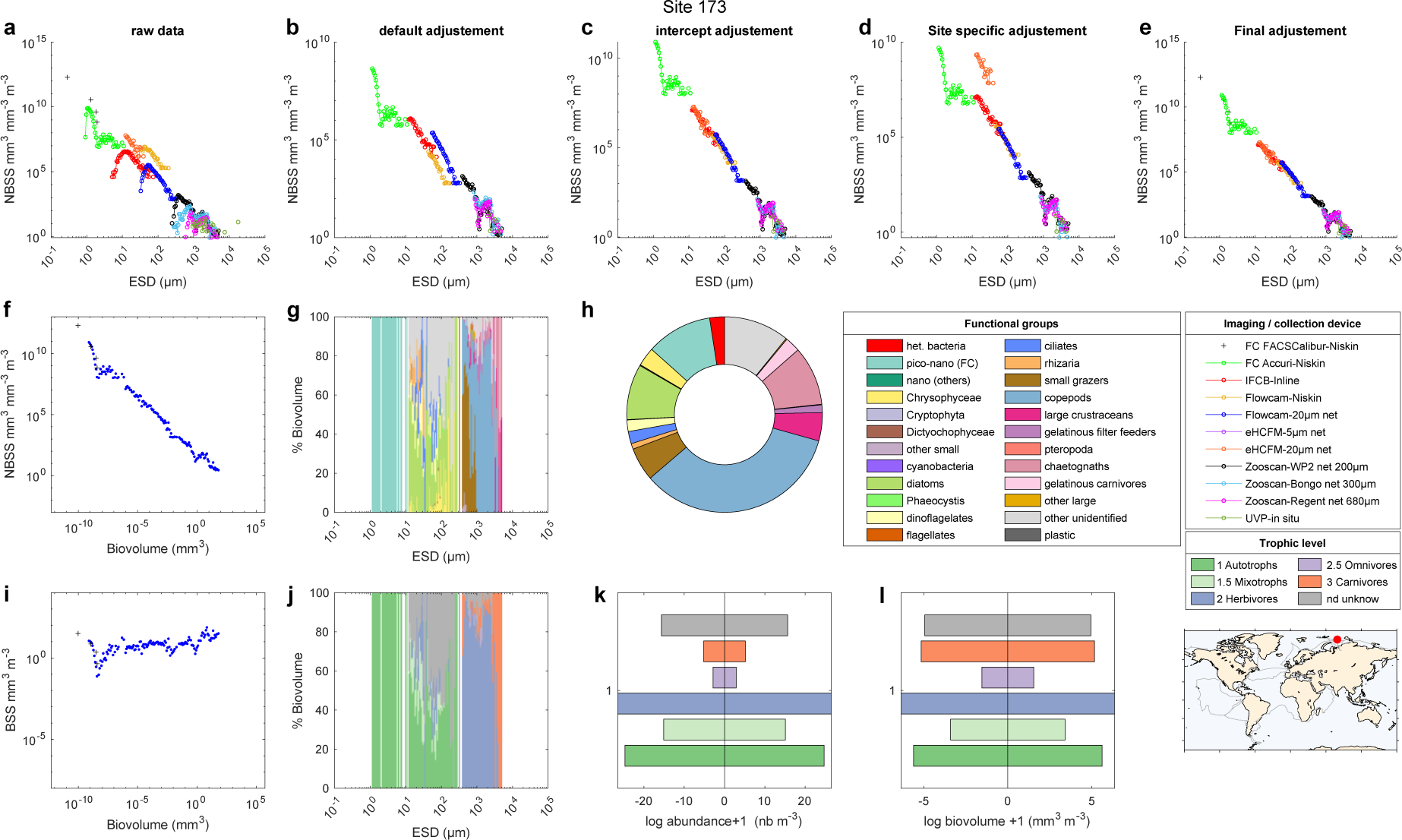
Meta-plankton assemblage example (Station 173). Living organisms raw NB- SS (a) and the different potential adjustments (b) default, (c) intercept, (d) site specific and (e) results of the final adjustment in NB-SS. Final assembled NB-SS (f) and B-SS (i) spectras and, size fractionated functional types proportions and trophic levels proportions (g,h) and total assemblage proportions as functional types (h) or trophic levels (k,l). Examples of all stations separated by day and night can be found as supplementary materials, together with version calculated in carbon units.

**Figure S8:**
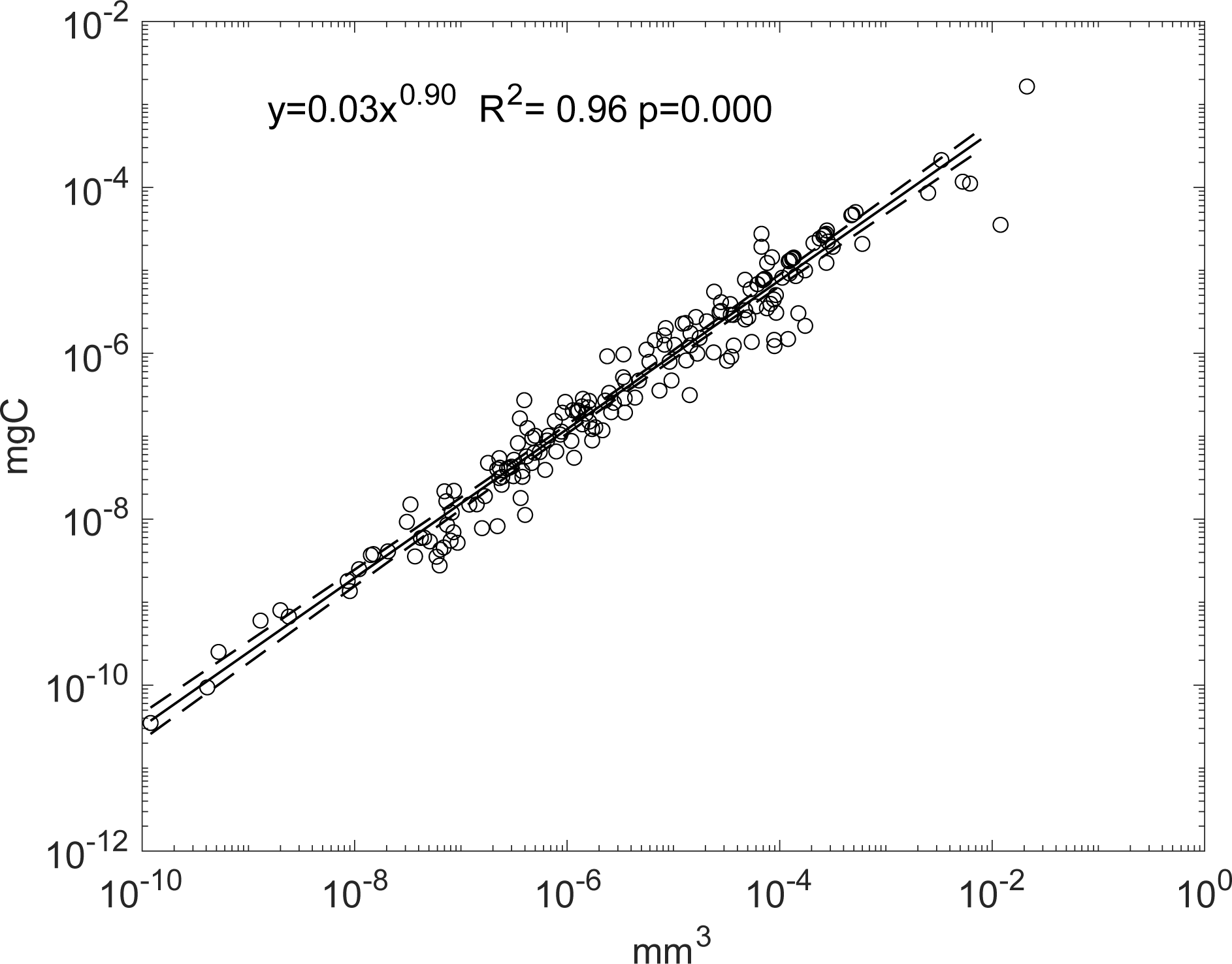
Relationship between cell biovolume and cell carbon in unicellular plankton organisms extracted from the bibliography (see Table SI-3).

